## Supplementary material for "MONET: Multi-omic patient module detection by omic selection": All supplementary information

### Simulations

#### Simulation I

We simulated samples from five modules, each with 60 samples, and two omics of dimension 500. Modules 1-3 cover both omics, module 4 only omic 1, and module 5 only omic 2. Each module has a *center* in each omic that it covers, and each sample is drawn from a normal multi-variate distribution around that center with unit covariance matrix. The center of each module was a vector of length 500 of all zeroes, except for 125 entries that equal 1 and characterize this module. The 125 module-specific features were disjoint for each module. The center of module 4 in omic 2 likewise used 125 entries that equal 1, but these entries were randomly sampled, and the covariance matrix of the normal distribution equaled $4 * I$, where $I$ is the identity matrix. Module 5 in omic 1 was generated in a similar manner. Five outlier samples were generated using multi-variate distribution with mean $\vec{0}$ and covariance $0.1 * I$.

#### Simulation II

We simulated samples from five modules, each with 30 samples, and three omics of dimension 500. As in simulation I, each module was simulated using multi-variate normal distribution in each omic that it covers. Module 1 had 100 non-zero entries in its center in omic 1. Modules 2-4 all had the same center in omic 1, with 100 non-zero entries. In omics 2 and 3, all modules had different centers, with different non-zero entries. In omic 2 each module had 20 non-zero entries, creating a weak clustering structure, and in omic 3 each had 40 non-zero entries.

### Datasets

#### Image dataset

The dataset was downloaded from: https://archive.ics.uci.edu/ml/machine-learning-databases/mfeat/ On 23 November 2018. It contains 2000 images of the digits 0-9, each digit with 200 images. Each image has the following 6 omics:

1 .mfeat-fou: 76 Fourier coefficients of the character shapes;
2 .mfeat-fac: 216 profile correlations;
3 .mfeat-kar: 64 Karhunen-Love coefficients;
4 .mfeat-pix: 240 pixel averages in 2 x 3 windows;
5 .mfeat-zer: 47 Zernike moments;
6. mfeat-mor: 6 morphological features.

#### scNMT

Data was downloaded from:
https://github.com/BIRSBiointegration/Hackathon/tree/master/scNMT-seq

#### Breast Invasive Carcinoma microarray

Agilent mRNA expression microarray data was downloaded from:
http://firebrowse.org/?cohort=BRCA&download_dialog=true

#### Benchmarked methods and software

Experiments were run on R version 3.5.2 64 bit.

For all methods, sequencing data (either RNA-seq or miRNA-seq) was log-transformed and features with 0 variance were removed. SNF, MONET and NEMO used all features. For MDI and clusternomics in each omic only the 2000 most highly variable features were kept. For BCC only 500 features were kept due to runtime. Each feature was then normalized to have mean 0 and standard deviation 1.

MONET – MONET was executed as described in the main text. In the image dataset all methods clustered the data into 10 clusters. MONET cannot get as input the number of clusters. Instead, when using NEMO to compute the edge weights, NEMO clustered each subsampled omic into 10 clusters, and the edge weights of each graph were shifted such that 10% of the weights were positive. This approach still does not guarantee that MONET finds the desired number of clusters.

To apply MONET to the scNMT data, we counted for every cell the number of promoters for which methylation was not measured. We removed the 25% of cells for which this number was lowest, and for every other cell we set to NA the methylation status for randomly selected promoters, such that all cells had the same number of measured promoters. We executed MONET as described before, where the number of clusters used by NEMO to compute the edge weights was 10.

To use MONET to discover gene modules, we only kept genes that were among the top 2000 most highly variable features in both the RNA-seq and microarray data. This left 1532 genes. Each gene was then normalized to have mean zero and standard deviation 1 in each omic. These omic matrices were then transposed and given as input to NEMO (which then normalized the *samples* to mean 0 and standard deviation 1, as it always does to its input).

SNF – SNF was executed as described previously in Rappoport et al., 2018[1].

NEMO – NEMO was executed as described previously in Rappoport et al., 2019[2]. Default parameters were used. When NEMO was run on single-cell data with many NA values per cell, as part of MONET's edge weight calculation, only values that are not NA were used to calculate distances between cells.

BCC – We used the R package bayesCC available here: https://github.com/ttriche/bayesCC. We executed BCC in parallel for K clusters with K ranging from 2 to 15. For each K, we executed the function bayesCC with IndivAlpha=TRUE and a maximum of 10,000 iterations. We selected the solution with highest mean adherence. Execution time reported is the wall clock time.

MDI –We used a recent implementation of MDI (mdipp-1.0.1) available here: https://warwick.ac.uk/fac/cross_fac/zeeman_institute/zeeman_research/software/. We executed the binary without CUDA support. To parse the output of the binary, determine the number of clusters, and perform the clustering, we used R scripts available here: https://github.com/cyversewarwick/mdipp/. Execution time reported only considers the data normalization, the execution time reported by the MDI binary, and the output parsing. The time taken to write to disk the input to the binary is not included.

Clusternomics – We used the R package clusternomics available here: https://github.com/evelinag/clusternomics. We executed clusternomics with either 3 or 5 clusters per omic, and with 5, 10, …, 30 global clusters, as performed in the authors' analysis. All these executions were done in parallel. For each of these options we called the function contextCluster with a maximum of 10,000 iterations, 3 iteration lag, 5000 iterations burn-in, and while modeling the data as normal with diagonal covariance matrix. We chose the clustering solution with minimal deviance information criterion. Execution time reported is the wall clock time.

#### Hardware

All experiments for timing the different methods were performed on a cluster:

Linux 4.9 72 CPUs, 2300 MHz each 756 GB RAM 64 bit operating system.
Since several methods use parallelization, the presented time for all methods is the wall clock time.

### Supplementary Tables

Supplementary Table 1 - TCGA log-rank p-values

| | nemo | snf | clusternomics | mdi | bcc | monet |
|---|---|---|---|---|---|---|
| aml | **7.3E-03** | **1.3E-03** | **3.3E-03** | **1.3E-02** | **3.9E-03** | **3.9E-02** |
| breast | **3.8E-02** | 9.9E-02 | 6.0E-02 | 1.0E-01 | **4.1E-03** | 3.9E-01 |
| colon | 6.5E-01 | 6.9E-01 | 5.5E-01 | 3.9E-01 | 8.8E-01 | 1.2E-01 |
| gbm | **1.3E-02** | **6.2E-05** | **1.5E-02** | **3.1E-03** | 2.6E-01 | **2.8E-04** |
| kidney | 6.8E-02 | **8.5E-03** | 1.6E-01 | **2.4E-02** | **4.9E-02** | **3.4E-02** |
| liver | **4.4E-04** | 6.6E-01 | 1.6E-01 | 5.4E-02 | 3.2E-01 | **1.1E-02** |
| lung | 4.3E-01 | 2.5E-01 | 4.1E-01 | **4.5E-02** | 9.2E-01 | 2.9E-01 |
| melanoma | **1.5E-04** | 2.4E-01 | **1.4E-02** | 8.2E-01 | 8.1E-01 | 7.4E-01 |
| ovarian | 7.1E-01 | 5.7E-01 | 5.8E-01 | 9.5E-01 | 9.1E-01 | **3.8E-02** |
| sarcoma | **1.5E-02** | **8.0E-03** | 6.8E-02 | **6.6E-03** | **6.0E-04** | **4.6E-02** |

Supplementary Table 2 - TCGA runtime (seconds)

| | nemo | snf | clusternomics | mdi | bcc | monet |
|---|---|---|---|---|---|---|
| aml | 8 | 9 | 12016 | 2633 | 32010 | 14 |
| breast | 28 | 55 | 37140 | 8928 | 90877 | 496 |
| colon | 16 | 16 | 19946 | 3305 | 37468 | 18 |
| gbm | 11 | 9 | 19262 | 3920 | 30146 | 46 |
| kidney | 12 | 8 | 17414 | 2940 | 21395 | 26 |
| liver | 19 | 20 | 33109 | 5511 | 53957 | 87 |
| lung | 15 | 17 | 28948 | 5125 | 51407 | 75 |
| melanoma | 19 | 29 | 41056 | 6802 | 64761 | 166 |
| ovarian | 10 | 13 | 28509 | 4191 | 46394 | 48 |
| sarcoma | 11 | 11 | 19041 | 3983 | 41910 | 59 |

Supplementary Table 3 - TCGA number of clusters

| | nemo | snf | clusternomics | mdi | bcc | monet |
|---|---|---|---|---|---|---|
| aml | 5 | 4 | 25 | 9 | 8 | 3 |
| breast | 3 | 2 | 30 | 11 | 3 | 6 |
| colon | 3 | 3 | 30 | 8 | 2 | 3 |
| gbm | 4 | 2 | 25 | 9 | 2 | 5 |
| kidney | 12 | 4 | 20 | 3 | 4 | 6 |
| liver | 5 | 2 | 25 | 10 | 7 | 5 |
| lung | 2 | 2 | 20 | 12 | 4 | 6 |
| melanoma | 5 | 3 | 30 | 8 | 3 | 6 |
| ovarian | 3 | 3 | 30 | 10 | 7 | 4 |
| sarcoma | 3 | 3 | 30 | 9 | 3 | 4 |

Supplementary Table 4 – venous invasion status for modules in MONET's solution on Ovarian Serous Cystadenocarcinoma data from TCGA

| MODULE | NO | YES |
|---|---|---|
| 1 | 15 | 3 |
| 2 | 8 | 24 |
| 3 | 3 | 5 |
| 4 | 10 | 17 |

Supplementary Table 5 - mutations tested for enrichment in MONET's solution on Ovarian Serous Cystadenocarcinoma data from TCGA

| gene | pval | qval | is_previously_known |
|---|---|---|---|
| TP53 | 0.57 | 1.00 | TRUE |
| BRCA1 | 0.40 | 1.00 | TRUE |
| CSMD3 | 0.14 | 1.00 | TRUE |
| NF1 | 0.45 | 1.00 | TRUE |
| CDK12 | 0.22 | 1.00 | TRUE |
| FAT3 | 0.15 | 1.00 | TRUE |
| GABRA6 | 0.16 | 1.00 | TRUE |
| BRCA2 | 0.27 | 1.00 | TRUE |
| RB1 | 0.78 | 1.00 | TRUE |
| TTN | 0.04 | 1.00 | FALSE |
| MUC16 | 0.25 | 1.00 | FALSE |
| HMCN1 | 0.78 | 1.00 | FALSE |
| USH2A | 0.88 | 1.00 | FALSE |
| AHNAK2 | 0.56 | 1.00 | FALSE |
| MUC17 | 0.54 | 1.00 | FALSE |
| DST | 0.27 | 1.00 | FALSE |
| RYR2 | 0.02 | 0.77 | FALSE |
| DNAH5 | 0.43 | 1.00 | FALSE |
| LRP2 | 0.73 | 1.00 | FALSE |
| APOB | 0.27 | 1.00 | FALSE |
| COL6A3 | 0.33 | 1.00 | FALSE |
| FAT1 | 0.78 | 1.00 | FALSE |
| HYDIN | 0.64 | 1.00 | FALSE |
| LRP1B | 0.20 | 1.00 | FALSE |
| PKHD1 | 0.41 | 1.00 | FALSE |
| ZNF614 | 0.41 | 1.00 | FALSE |
| AHNAK | 0.21 | 1.00 | FALSE |
| DNAH3 | 0.40 | 1.00 | FALSE |
| FLG | 0.54 | 1.00 | FALSE |
| CACNA1C | 0.80 | 1.00 | FALSE |
| COL22A1 | 0.26 | 1.00 | FALSE |
| GPR98 | 0.53 | 1.00 | FALSE |
| HUWE1 | 0.66 | 1.00 | FALSE |

Supplementary Table 6 – highly expressed genes in Module 3 of MONET's solution on Ovarian cancer.

| name | pval | qval |
|---|---|---|
| hsa.let.7g | 5.2E-06 | 3.2E-03 |
| hsa.mir.101.1 | 4.7E-05 | 2.9E-02 |
| hsa.mir.17 | 1.1E-07 | 6.8E-05 |
| hsa.mir.181c | 5.8E-05 | 3.6E-02 |
| hsa.mir.181d | 5.7E-06 | 3.5E-03 |
| hsa.mir.19b.1 | 1.6E-07 | 9.8E-05 |
| hsa.mir.200b | 2.5E-06 | 1.5E-03 |
| hsa.mir.202 | 3.0E-05 | 1.9E-02 |
| hsa.mir.20a | 3.3E-05 | 2.0E-02 |
| hsa.mir.26a.2 | 1.6E-05 | 9.6E-03 |
| hsa.mir.30c.2 | 3.1E-05 | 1.9E-02 |
| hsa.mir.506 | 1.3E-05 | 7.9E-03 |
| hsa.mir.508 | 3.1E-06 | 1.9E-03 |
| hsa.mir.509.1 | 1.1E-05 | 6.9E-03 |
| hsa.mir.509.2 | 8.4E-06 | 5.2E-03 |
| hsa.mir.509.3 | 9.4E-06 | 5.8E-03 |
| hsa.mir.514.1 | 2.4E-05 | 1.5E-02 |
| hsa.mir.514.2 | 2.7E-05 | 1.6E-02 |
| hsa.mir.514.3 | 1.8E-05 | 1.1E-02 |
| hsa.mir.99a | 1.3E-07 | 7.7E-05 |

Supplementary Table 7 – Mouse embryonic day of development for MONET's solution on scNMT data.

| | 1 | 2 | 3 | 4 | 5 | 6 | 7 | 8 | 9 | 10 | 11 |
|---|---|---|---|---|---|---|---|---|---|---|---|
| E4.5 | 42 | 0 | 23 | 0 | 0 | 0 | 0 | 0 | 0 | 0 | 0 |
| E5.5 | 0 | 1 | 0 | 0 | 11 | 0 | 4 | 0 | 66 | 0 | 0 |
| E6.5 | 0 | 6 | 0 | 0 | 18 | 88 | 67 | 1 | 1 | 0 | 0 |
| E7.5 | 0 | 3 | 0 | 42 | 0 | 58 | 0 | 33 | 0 | 57 | 18 |

Supplementary Table 8 – Cell types for MONET's solution on scNMT data.

| | 1 | 2 | 3 | 4 | 5 | 6 | 7 | 8 | 9 | 10 | 11 |
|---|---|---|---|---|---|---|---|---|---|---|---|
| Ectoderm | 0 | 0 | 0 | 0 | 0 | 29 | 0 | 0 | 0 | 5 | 0 |
| Endoderm | 0 | 3 | 0 | 0 | 0 | 0 | 0 | 0 | 0 | 13 | 18 |
| Epiblast | 42 | 0 | 0 | 0 | 0 | 98 | 46 | 0 | 67 | 3 | 0 |
| ExE_ectoderm | 0 | 0 | 0 | 0 | 0 | 0 | 4 | 0 | 0 | 0 | 0 |
| Mesoderm | 0 | 0 | 0 | 41 | 0 | 1 | 5 | 34 | 0 | 23 | 0 |
| Primitive_endoderm | 0 | 0 | 23 | 0 | 0 | 0 | 0 | 0 | 0 | 0 | 0 |
| Primitive_Streak | 0 | 0 | 0 | 1 | 0 | 18 | 15 | 0 | 0 | 12 | 0 |
| Visceral_endoderm | 0 | 7 | 0 | 0 | 29 | 0 | 0 | 0 | 0 | 0 | 0 |

Supplementary Table 9 – Mouse embryonic day of development and cell type for MONET's solution on scNMT data.

| | 1 | 2 | 3 | 4 | 5 | 6 | 7 | 8 | 9 | 10 | 11 |
|---|---|---|---|---|---|---|---|---|---|---|---|
| E4.5_Epiblast | 42 | 0 | 0 | 0 | 0 | 0 | 0 | 0 | 0 | 0 | 0 |
| E4.5_NA | 0 | 0 | 0 | 0 | 0 | 0 | 0 | 0 | 0 | 0 | 0 |
| E4.5_Primitive_endoderm | 0 | 0 | 23 | 0 | 0 | 0 | 0 | 0 | 0 | 0 | 0 |
| E5.5_Epiblast | 0 | 0 | 0 | 0 | 0 | 0 | 4 | 0 | 66 | 0 | 0 |
| E5.5_Visceral_endoderm | 0 | 1 | 0 | 0 | 11 | 0 | 0 | 0 | 0 | 0 | 0 |
| E6.5_Epiblast | 0 | 0 | 0 | 0 | 0 | 71 | 42 | 0 | 1 | 0 | 0 |
| E6.5_ExE_ectoderm | 0 | 0 | 0 | 0 | 0 | 0 | 4 | 0 | 0 | 0 | 0 |
| E6.5_Mesoderm | 0 | 0 | 0 | 0 | 0 | 1 | 5 | 1 | 0 | 0 | 0 |
| E6.5_NA | 0 | 0 | 0 | 0 | 0 | 0 | 1 | 0 | 0 | 0 | 0 |
| E6.5_Primitive_Streak | 0 | 0 | 0 | 0 | 0 | 16 | 15 | 0 | 0 | 0 | 0 |
| E6.5_Visceral_endoderm | 0 | 6 | 0 | 0 | 18 | 0 | 0 | 0 | 0 | 0 | 0 |
| E7.5_Ectoderm | 0 | 0 | 0 | 0 | 0 | 29 | 0 | 0 | 0 | 5 | 0 |
| E7.5_Endoderm | 0 | 3 | 0 | 0 | 0 | 0 | 0 | 0 | 0 | 13 | 18 |
| E7.5_Epiblast | 0 | 0 | 0 | 0 | 0 | 27 | 0 | 0 | 0 | 3 | 0 |
| E7.5_Mesoderm | 0 | 0 | 0 | 41 | 0 | 0 | 0 | 33 | 0 | 23 | 0 |
| E7.5_NA | 0 | 0 | 0 | 0 | 0 | 0 | 0 | 0 | 0 | 1 | 0 |
| E7.5_Primitive_Streak | 0 | 0 | 0 | 1 | 0 | 2 | 0 | 0 | 0 | 12 | 0 |

### Supplementary Figures

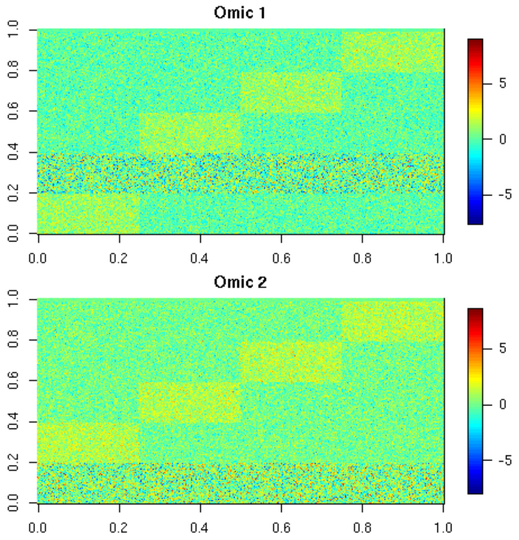

Supplementary Figure 1 – raw data for simulation I. Rows are samples and columns are features.

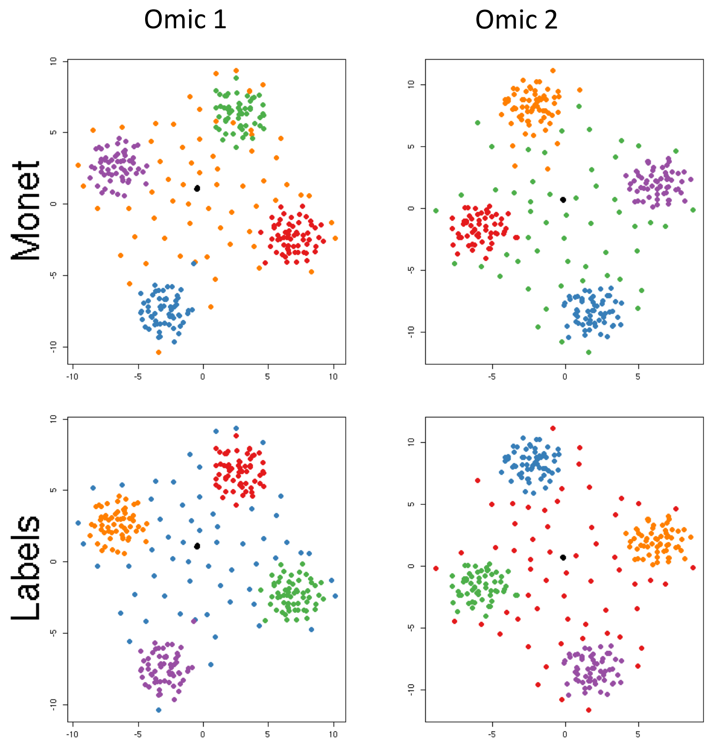

Supplemenary Figure 2 – t-sne visualization of the raw data in simulation I. Samples are colored by MONET's output (top) or the true labeling (bottom). Lonely samples are colored in black.

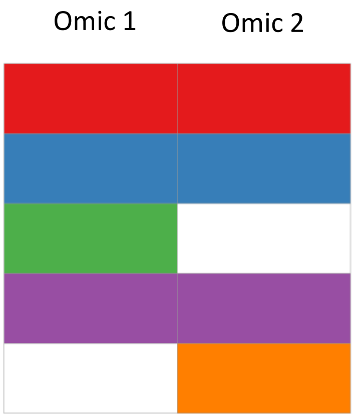

Supplementary Figure 3 – Module omics identified by MONET on simulation I. Each row signifies a module and each column an omic. Colored panels indicate that the omic is covered by the module, white indicates that it is not.

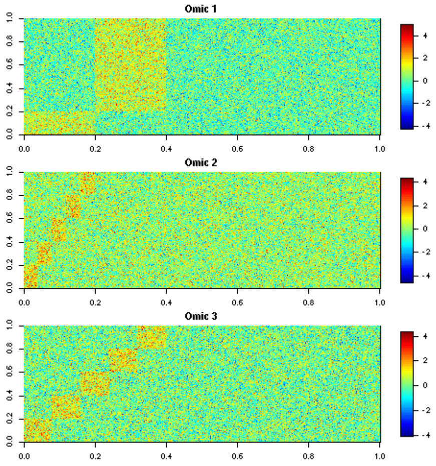

Supplementary Figure 4 – raw data for simulation II. Rows are samples and columns are featu[illegible]

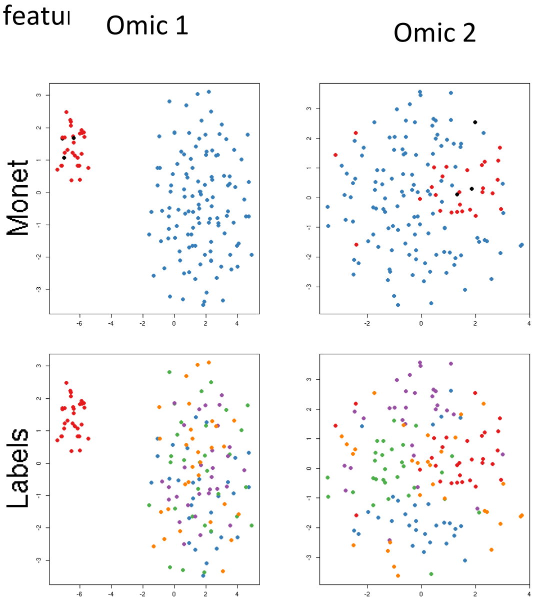

Supplementary Figure 5 – t-sne visualization of the raw data in simulation II using only the first 2 omics. Samples are colored by MONET's output (top) or the true labeling (bottom).

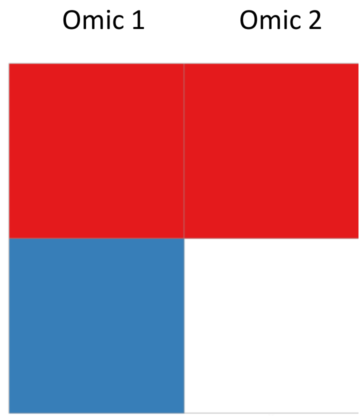

Supplementary Figure 6 – omics covered by each MONET module in simulation II when using only the first two omics.

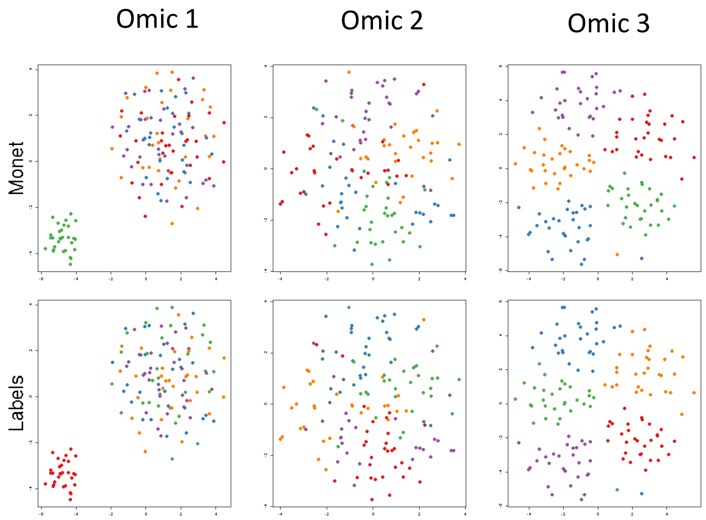

Supplementary Figure 7 – t-sne of raw data in simulation II using all 3 omics. Samples are colored by MONET's output (top) or the true labeling (bottom).

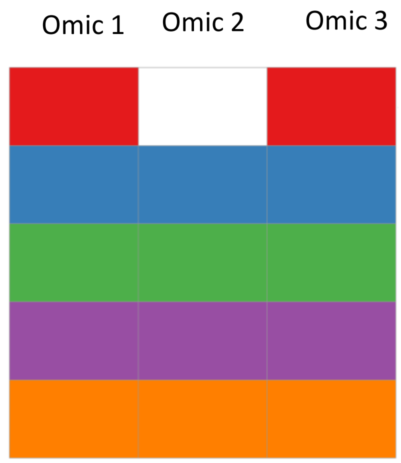

Supplementary Figure 8 – Omics covered by each MONET module in simulation II when using all the three omics.

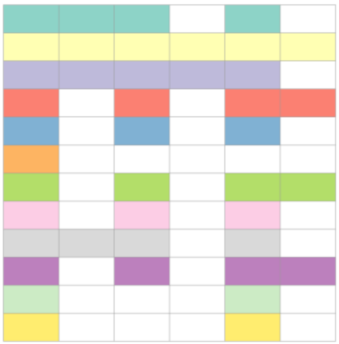

Supplementary Figure 9 – Omics covered by each MONET module in the image dataset. Columns are omics and rows are modules.

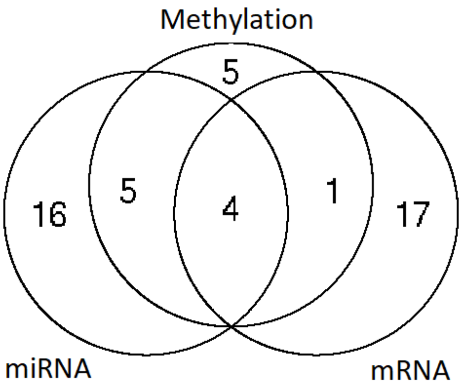

Supplementary Figure 10 – Distribution of omics covered by MONET modules across all 10 TCGA cancer subtypes.

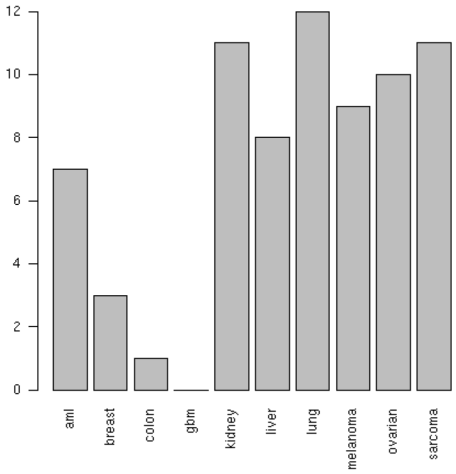

Supplementary Figure 11 – Number of outliers (lonely samples) reported by MONET per cancer type.

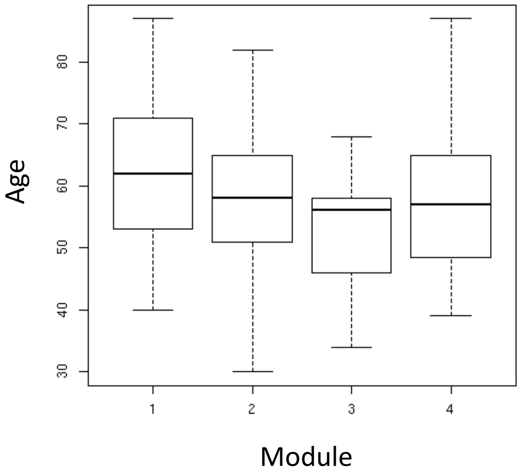

Supplementary Figure 12 – Age distribution across 4 MONET modules for Ovarian cancer.

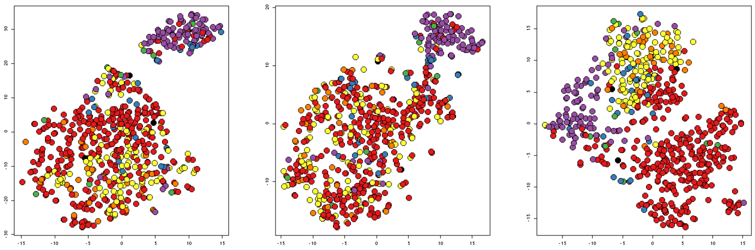

Supplementary Figure 13 – t-sne visualization of MONET's solution on Breast cancer from TCGA. Samples are colored by MONET's output (top) or the true labeling (bottom).

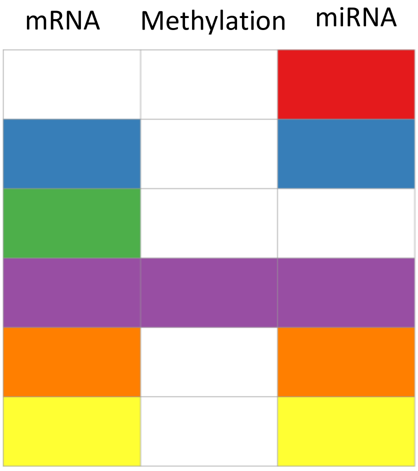

Supplementary Figure 14 – Omics covered by each MONET module in Breast cancer data from TCGA. Columns are omics and rows are modules.

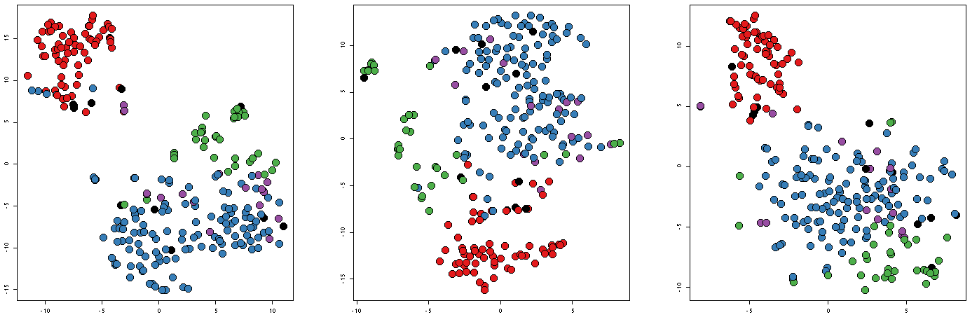

Supplementary Figure 15 – t-sne visualization of MONET's solution on Sarcoma data from TCGA. Samples are colored by MONET's output (top) or the true labeling (bottom).

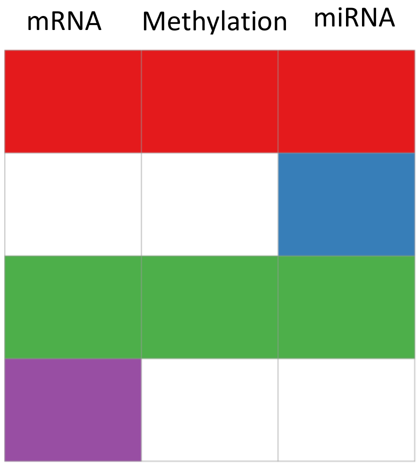

Supplementary Figure 16 – Omics covered by each MONET module in Sarcoma data from TCGA. Columns are omics and rows are modules.

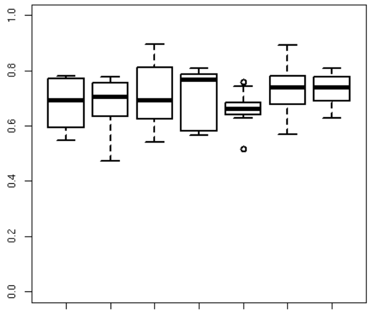

Supplementary Figure 17 – ARI in partial datasets experiments for the image dataset. Shown is the ARI distribution for samples dropped in each omic, and for the whole dataset (rightmost box), compared to the ground truth solution.

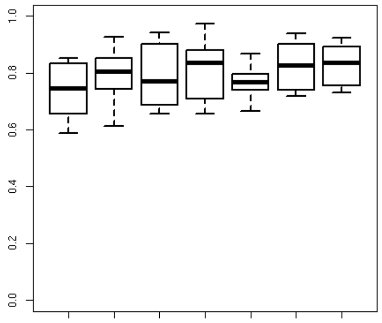

Supplementary Figure 18 – same as Supp Fig 12, only comparing to the solution on all samples.

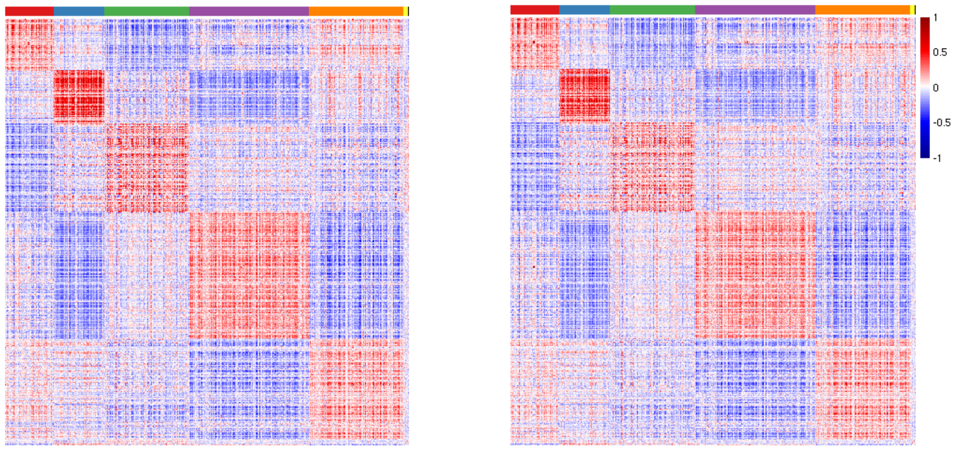

Supplementary Figure 19 – Correlation matrix of genes in the Breast Invasive Carcinoma dataset using RNA-seq (left) and microarrays (right). Genes are marked by their MONET module.
